# miRador: a fast and precise tool for the prediction of plant miRNAs

**DOI:** 10.1101/2021.03.24.436803

**Authors:** Reza K. Hammond, Pallavi Gupta, Parth Patel, Blake C. Meyers

**Author notes:** **Correspondence to:** (B.C.M.).

## Abstract

Plant microRNAs (miRNAs) are short, non-coding RNA molecules that restrict gene expression via post-transcriptional regulation and function in several essential pathways including development, growth, and stress responses. Accurately identifying miRNAs in populations of small RNA (sRNA) sequencing libraries is a computationally intensive process which has resulted in the misidentification of inaccurately annotated miRNA sequences. In recent years, criteria for miRNA annotation have been refined to reduce these misannotations. Here, we describe *miRador*, a novel miRNA identification tool that utilizes the most up-to-date, community-established criteria for accurate identification of miRNAs in plants. We combine target prediction and Parallel Analysis of RNA Ends (PARE) data to assess the precision of the miRNAs identified by *miRador*. We compare *miRador* to other commonly used miRNA prediction tools and we find that *miRador* is at least as precise as other prediction tools while being significantly faster than other tools.

## INTRODUCTION

Eukaryotic genomes have evolved to encode diverse classes of small non-coding RNA (sRNA) molecules that function in partially overlapping epigenetic silencing pathways. It is believed that sRNAs evolved as a means of defense against RNA viral infections and for silencing transposable elements, and that they later adapted to regulate the expression of endogenous genes (Borges & Martienssen, 2015; Chen et al., 2018). In plants, microRNAs (miRNAs) are a subclass of sRNA that function to regulate gene expression via posttranscriptional gene silencing, operating in several pathways important to plants, including development, growth, and stress responses. The miRNA biogenesis and miRNA-induced silencing pathways have been extensively studied in *Arabidopsis*, and, to date, these pathways have been found to be well conserved in all land plants that have been examined.

Given the significance of miRNAs in gene regulation, a set of standards were created for the accurate annotation of miRNAs in the organisms in which they were identified. The first effort to define standards for miRNA annotation was published in 2003 and primarily relied on a combination of evidence of both expression and biogenesis (Ambros et al., 2003); criteria for plant miRNA annotation were not explicitly defined, separate from animals. Evidence of expression included, at that time, the accumulation of candidate miRNA in gel blots and the identification of a candidate miRNA in a library of cDNAs made from size-fractionated RNAs. Evidence of biogenesis included the prediction of a fold-back miRNA precursor and the mature sequence mapping entirely to a single arm of that hairpin, conservation of the candidate miRNA and its predicted precursor secondary structure, and the detection of increased precursor accumulation in *dicer* mutants. By today’s standards, it is understood that these requirements alone are insufficient to properly classify miRNAs. In particular, the accumulation of a candidate miRNA does not differentiate it from the other many classes of sRNAs; conservation with a known miRNA assumes that the sequence is required to be a miRNA in the newly studied organism and that the original annotation was correct; in addition, reduced accumulation in *dicer* mutants ignores the partial redundancy of plant *DCL* genes (Axtell & Meyers, 2018).

In 2008, another community effort was made to redefine miRNA annotation standards, at that point integrating observations from then-new, deep sequencing technologies in order to reduce false-positive miRNA annotations. These criteria required for validation of a candidate miRNA that the miRNA:miRNA* duplex is identified on a hairpin precursor (Axtell & Meyers, 2018; Meyers et al., 2008). Those requirements were summarized into eight specific rules that largely served as the basis of numerous first-generation plant miRNA prediction tools. Subsequently, in 2018, these rules were updated to reflect the changes in understanding of plant miRNAs and their biogenesis that resulted from the massive amounts of data that had accumulated over a decade’s worth of sequencing, genomics, and analysis (**Table 1**). The motivation for making changes to the criteria was to leverage the increased understanding of miRNA biogenesis to further minimize false positive miRNA annotations, by employing stricter annotation requirements for candidate miRNAs. This update did more than just make the requirements stricter, however; some rules were relaxed to prevent false negatives in miRNA predictions (Axtell & Meyers, 2018).

**Table 1:** Past and current criteria for plant miRNA annotations.

|  | **2008 Criteria** | **2018 Criteria** |
| --- | --- | --- |
| 1 | One or more miRNA:miRNA* duplexes with two-nucleotide 3' overhangs | Add requirements that exclude secondary stems or large loops (larger than 5 nucleotides) in the miRNA:miRNA* duplex and limit precursor length to 300 nucleotides |
| 2 | Confirmation of both the mature miRNA and its miRNA* | Disallow confirmation by blot; sRNA-seq only |
| 3 | miRNA:miRNA* duplex contains $\leq 4$ mismatched bases | Up to five mismatched positions, only three of which are nucleotides in asymmetric bulges |
| 4 | The duplex has at most one asymmetric bulge containing at most two bulged nucleotides | Up to five mismatched positions, only three of which are nucleotides in asymmetric bulges |
| 5 | $\geq 75\%$ of reads from exact miRNA or miRNA* | Include one-nucleotide positional variants of miRNA and miRNA* when calculating precision |
| 6 | Replication suggested but not required | Required; novel annotations should meet all criteria in at least two sRNA-seq libraries (biological replicates) |
| 7 | Homologs, orthologs, and paralogs can be annotated without expression data, provided all criteria met for at least one locus in at least one species | Homology-based annotations should be noted as provisional, pending actual fulfillment of all criteria by sRNA-seq |
| 8 | miRNA length not an explicit consideration | No RNAs $< 20$ nucleotide or $> 24$ nucleotides should be annotated as miRNAs. Annotations of 23- or 24-nucleotide miRNAs require extremely strong evidence. |

Note: 2008 criteria from Meyers et al., 2008; 2018 criteria from Axtell & Meyers, 2018.

With the development of these new rules, however, it became imperative to develop a computational tool to improve plant miRNA predictions by implementing and enforcing the rules. For this reason, we developed a novel plant miRNA prediction tool, *miRador*, that utilizes the updated rules to create what we assert is one of the most precise plant miRNA prediction tools available today. In comparison to more commonly-used plant miRNA prediction tools, we found that *miRador* is faster and at least as precise as existing plant miRNA prediction tools. We also developed new functionality within *miRador* to provide users with important conservation insights of predicted miRNAs that other tools lack. In conjunction with *sPARTA*, a target prediction and validation tool (Kakrana et al., 2014), we found that we can generate high quality miRNA predictions in a variety of plant species.

## RESULTS

### Description of the miRador miRNA prediction strategy

A flowchart depicting the pipeline is depicted in **Figure 1**. Upon initiating a miRNA prediction run with *miRador*, the application utilizes the user-provided genome file to identify inverted repeats within each chromosome using *einverted* (Rice et al., 2000). *einverted* has several scoring parameters which can be manually set by the user, or the user can choose from three preset *miRador* options to generate a series of inverted repeats. These inverted repeats will serve as a base set of candidate precursor miRNAs and are stored into a Python dictionary for subsequent analysis.

**Figure 1:**
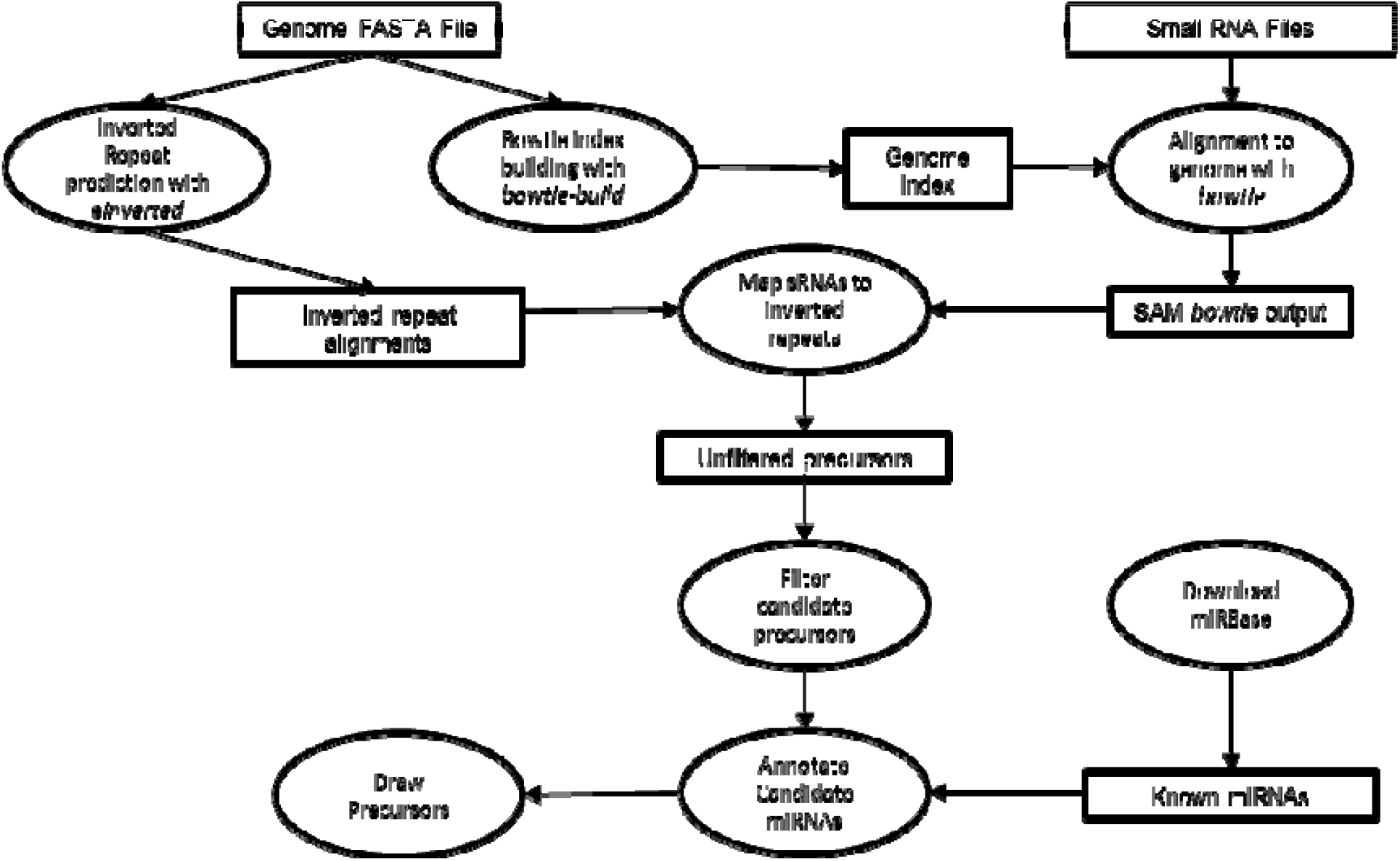
Pipeline of *miRador.* *miRador* requires two sets of input files: 1) a genome FASTA file, and 2) sequenced small RNA files. Processing begins with the prediction of inverted repeats on each chromosome with *einverted.* Small RNA sequences are subsequently mapped to the genome, for alignment against inverted repeats, which are the initial set of “candidate precursor miRNAs.” These are then filtered utilizing the 2018 plant miRNA annotation criteria. Finally, these candidate miRNA**s** are annotated utilizing known plant miRNAs from miRBase, and replication requirements are enforced if a candidate miRNA is novel.

Small RNA libraries, which can be provided as FASTA, FASTQ, or tag count (i.e. unique sequence and their read count) files, are independently processed from start to finish. Prior to mapping a library to a genome, sRNA sequences are read into another Python dictionary with sRNA sequences as keys and their read count as the attached value. This dictionary is then used to create a FASTA file containing only distinct (different) reads from the library. As this file contains only distinct reads, mapping time is significantly reduced as the process of mapping of each sRNA read to the genome occurs only once for a distinct sequence. Libraries are mapped to the genome with *bowtie* v1 and output in SAM format for immediate processing (Langmead et al., 2009).

This mapped file is parsed into nested Python dictionaries with sRNA positional coordinates as keys and a list of sequences that map to that position as the attached value. The use of dictionaries (Python’s built-in implementation of hash tables) is critical due to the data structure’s speed at accessing stored data when the index of the stored value is not known. Unlike lists, dictionaries can be queried on a string of characters, including numbers, to find their stored value. The average complexity of a search for a value in an array is O(n), whereas the average complexity of a search of a value in a hash table is O(1). The real-world impact of this design is reflected in the lookup times for cumulative millions of mapped read locations for which sRNAs need to be processed for mapping to inverted repeats. The mapping process is completed with the normalization of reads as reads per million (RPM).

For an inverted repeat to be considered a candidate precursor, a small RNA must map to both arms of the precursor. Thus, *miRador* iterates through each position on each arm of the inverted repeat to identify any sRNA that lies, in its entirety, on an arm of the inverted repeat. The previously created dictionaries are utilized to identify all sRNAs that map to each inverted repeat. Under the current criteria of plant miRNA annotation, no miRNA can be confirmed without a corresponding miRNA* (where “miRNA*”, read as miRNA-star, is the complement in the processed duplex). Thus, any inverted repeat without an sRNA mapping to both arms is immediately removed from the analysis. The remaining inverted repeats are analyzed to identify two sequences on opposite arms that could complete a miRNA:miRNA* duplex at this precursor. The criteria for a miRNA:miRNA* duplex are the following: (1) 2-nt 3’ overhangs on the alignment of the candidate miRNA and miRNA*, (2) up to 5 mismatched positions, only 3 of which may be nucleotides in asymmetric bulges (G-U pairing assessed ½ a mismatch), and (3) at least 75% of read abundance mapping to the precursor miRNA are from 1-nt positional variants of miRNA and miRNA*. *miRador* then assesses the alignment parameters by identifying the sRNA sequences on the predicted inverted repeats. If the two alignment criteria are met, then each 1-nt positional variants of both the candidate miRNA and the candidate miRNA* are identified to pool their abundances. If these abundances exceed 75% of the read abundance mapping to this inverted repeat, and the candidate miRNA has an abundance of at least 3 RPM per hit to the genome, then the miRNA:miRNA* duplex will be classified as a candidate miRNA within the library and will be analyzed further in the final step.

Upon the completion of prediction for each library, each confirmed miRNA:miRNA* duplex on the precursor miRNA is drawn utilizing *RNAFold* (Kerpedjiev et al., 2015). The alignments generated by *RNAFold* can differ from the *einverted* alignment due to differences in the scoring systems of the two tools.

Among the novel features of *miRador* is its annotation component that classifies candidate miRNAs. There are five classifications to which a candidate miRNA might be assigned: (1) known, (2) identical to known, (3) new member of existing family, (4) conserved outside the species of study, and (5) novel. When *miRador* initiates, it will automatically download all plant miRNAs that have been annotated in the selected version of miRBase, if it has not been downloaded already. In the annotation step, each candidate miRNA is analyzed for sequence similarity to any known miRNA via Basic Local Alignment Search Tool (BLAST) (Altschul et al., 1990; Altschul et al., 1997; Camacho et al., 2009). If the species being analyzed exists in miRBase, the General Feature Format (GFF) file is downloaded to determine the locations at which miRNAs and their precursors have been previously identified. A miRNA identified by *miRador* can only be classified as “known” if the miRNA was identified at the same position that it exists in miRBase. If the sequence is identical to a known miRNA, but not at the same location of any known miRNA, it is classified as “identical to a known miRNA”. A sequence is classified as a “new member of an existing family” if there are 5 or fewer differences to any known miRNA in the organism of study (a difference is referred to as a gap, mismatch, or bulge, while a G-U wobble is given a half point penalty). BLAST analysis may not find a similar miRNA within the organism of study, but it might identify a match in another plant. If a miRNA is identified as having 5 or fewer differences to any known miRNA outside of this organism, it will be classified as “conserved outside the organism of study”. This classification and “new member of an existing family” within the organism of study are not mutually exclusive, though the classification as a new member within the organism is given precedence and will appear first in the output file. Finally, if none of these classifications fit the candidate miRNA, it will be classified as “novel”. Given the novelty of this function and its general lack of dependency on *miRador*’s execution structure, we made this function available to run as a standalone tool to be used with the results of two of the most used miRNA prediction tools, *ShortStack* and *miRDeep-P2*. This tool is available in a Github repository https://github.com/rkweku/mirnaAnnotation.

The final step of *miRador* filters candidate miRNAs by ensuring novel miRNA families are predicted independently in multiple sRNA libraries. Upon assignment of classification of a candidate miRNA in the annotation step, *miRador* will determine the number of libraries in which it was predicted, if the candidate miRNA was not conserved within the organism of study. If the number of libraries predicting this candidate miRNA both exceeds one and exceeds 10% of the number of libraries provided for prediction, the candidate miRNA is confirmed by *miRador*. By requiring a candidate miRNA to be present in at least 10% of libraries, we ensure that *miRador* does not predict false positives when operating with numerous libraries. These final predictions are output to a CSV file for subsequent analysis by users.

### Assessing the predictive capabilities of miRador

miRBase is a repository for miRNA annotations that have been identified in the peer-reviewed literature (Kozomara et al., 2018), though it is important to note that it is not a gatekeeper in enforcing the quality of miRNA annotations (Axtell & Meyers, 2018). Outdated verification criteria and poor-quality sequencing libraries have both been implicated as a cause for improper verification of miRNAs within miRBase (Ludwig et al., 2017; Taylor et al., 2014). Despite these limitations of miRBase, previous miRNA prediction tools have used miRBase miRNAs as a source of true positive miRNAs (Johnson et al., 2016; Kuang et al., 2018). We, however, opted to determine the validity of a candidate miRNA as dictated through evidence of PARE (Parallel Analysis of RNA Ends) libraries (German et al., 2009), to determine a better set of true positives within each prediction set. This method of identifying true miRNAs has the added benefit of assessing the predictability of novel miRNAs, a useful feature given that these prediction tools exist largely to identify novel miRNAs. We determined evidence of miRNA-facilitated cleavage of mRNA via target prediction and PARE validation with *sPARTA* (Kakrana et al., 2014).

We first predicted miRNAs across 21 individual *Arabidopsis* seedling and flower libraries. *miRador* identified 228 total miRNAs, 131 of which were already present in miRBase (**Table 2**). One miRNA was found to be identical to an already known miRNA but at a different position of the genome, 45 were classified as new members of existing miRNA families, 19 were classified as conserved miRNAs that are known in other organisms, and 32 were classified as entirely novel. We utilized *sPARTA* to predict targets for each of the candidate miRNAs and validated their activity in 7 PARE libraries. Among the 228 candidate miRNAs, 179 had predicted targets with significant PARE activity (*P* < 0.05) at the predicted site of cleavage resulting in precision of 0.785.

**Table 2.**
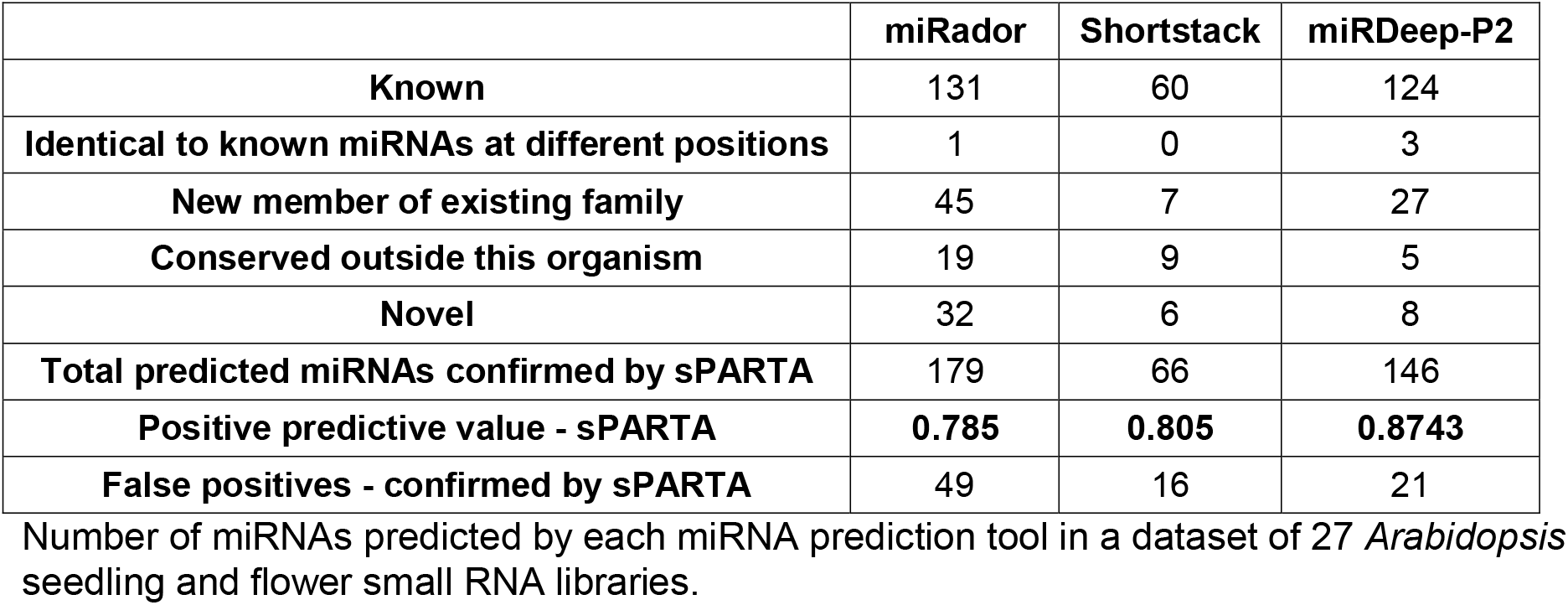
*Arabidopsis* miRNAs predicted by each tool.

We next extended our analysis to 30 rice libraries. From this data, *miRador* identified 391 candidate miRNAs, 357 of which had significant PARE activity at their predicted sites of cleavage in four PARE panicle libraries resulting in a total precision of 0.913 (**Table 3**). This precision exceeded that of the *Arabidopsis* miRNAs, which may largely be attributable to these sRNA and PARE libraries being generated from the same amount of isolated, low-input material. Of the 391 total predictions, the vast majority were classified as novel miRNAs. While 76 miRNAs that were identified are already known, 252 were novel.

**Table 3.** Rice miRNAs predicted by each tool.

|  | **miRador** | **Shortstack** | **miRDeep-P2** |
| --- | --- | --- | --- |
| **Known** | 76 | 38 | 111 |
| **Identical to known miRNAs at different positions** | 2 | 0 | 3 |
| **New member of existing family** | 44 | 7 | 141 |
| **Conserved outside this organism** | 17 | 1 | 6 |
| **Novel** | 252 | 38 | 178 |
| **Total predicted miRNAs confirmed by sPARTA** | 357 | 73 | 403 |
| **Positive predictive value - sPARTA** | **0.913** | **0.869** | **0.918** |
| **False positives - confirmed by sPARTA** | 34 | 11 | 36 |

Number of miRNAs predicted by each miRNA prediction tool in a dataset of 12 rice anther small RNA libraries.

Subsequent predictions with 44 maize anther, seedling, and tassel libraries identified 173 maize miRNAs. Among these candidate miRNAs, 149 had predicted targets with significant PARE activity at the predicted site of cleavage resulting in precision of 0.861 (**Table 4**). *miRador* identified few miRNAs that were novel to maize (16 total). However, nearly half of the total predictions made by *miRador* were known mature miRNA sequences derived from novel *MIRNA* genes (16) as well as new members of known miRNA families (60). Unlike the *Arabidopsis* and rice predictions, there were very few novel predictions with only 9 predicted miRNAs being novel. Overall, the results of these analyses indicate that *miRador* can predict miRNAs from an assortment of sRNA sequencing files from multiple organisms with a high degree of precision, as determined by PARE activity at the predicted sites of miRNA action.

**Table 4.**
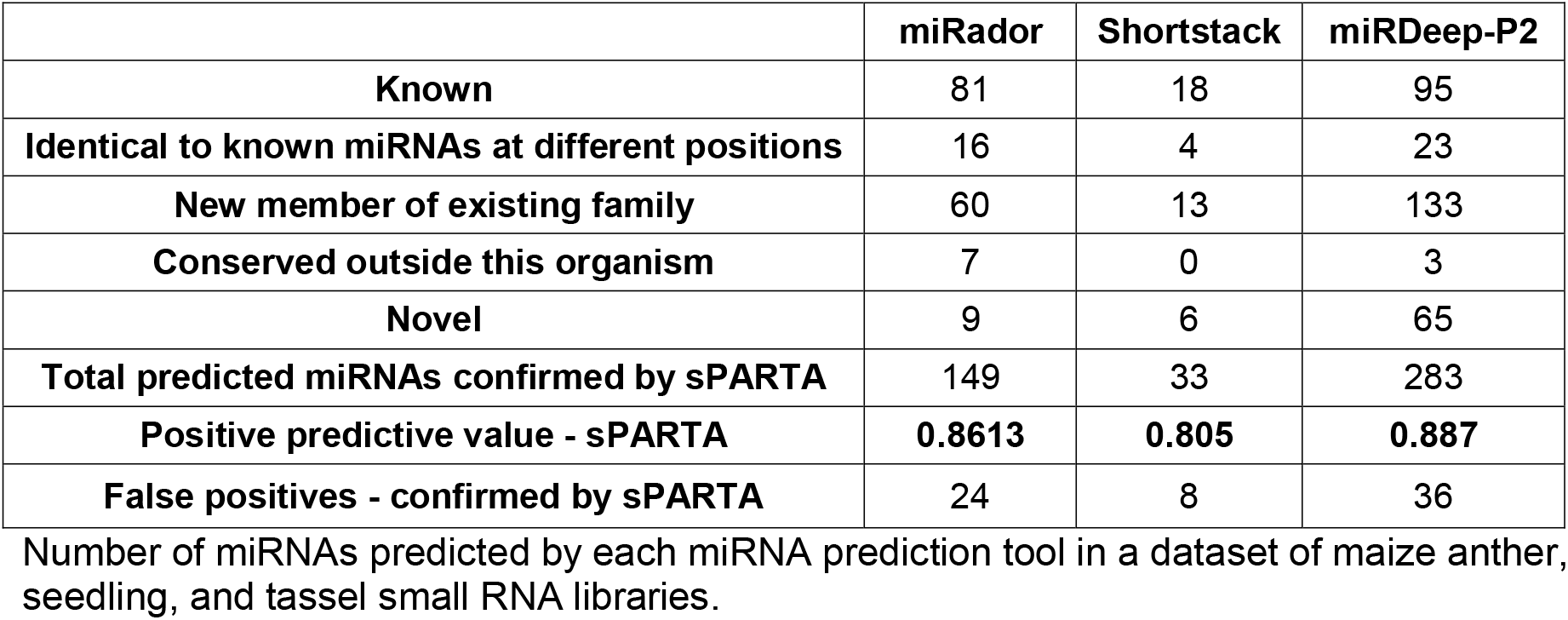
Maize miRNAs predicted by each tool.

### Comparison to other miRNA prediction tools

To better assess the value of *miRador*’s predictive capabilities, we compared its performance with those of two commonly used plant miRNA prediction tools: *ShortStack* and *miRDeep-P2. ShortStack* (Johnson et al., 2016) is a comprehensive analytical tool that classifies mapped sRNA sequencing data. *miRDeep-P2* is exclusively a plant miRNA prediction tool, and was published as an update to the popular *miRDeep-P (Kuang et al., 2018)*. We used each of the three prediction tools to predict miRNAs in the same *Arabidopsis*, rice, and maize libraries described above. *miRDeep-P2* can only perform predictions on single libraries, so its predictions across multiple runs were merged to create a single set of predicted miRNAs for all libraries within a dataset. Additionally, *miRDeep-P2* utilizes an abundance cutoff, but it bypasses this cutoff if the candidate miRNA differs by up to 1 nucleotide from any miRBase miRNA. Since we wish to address the merits of each of these prediction tools using their default behavior for all candidate miRNAs, we removed this bypass in our assessment. *ShortStack* and *miRDeep-P2* miRNA predictions were also annotated using the miRNA annotation component from *miRador* to consistently annotate the predicted miRNAs of each tool. This approach also has the added benefit of enforcing consistent replication requirements for *miRDeep-P2*’s predictions as there is no utility for this built into *miRDeep-P2*, due to its capability to only process single libraries.

We found that *miRador* and *miRDeep-P2* both identified a similar number of known *Arabidopsis* miRNAs, 131 and 124, respectively while *ShortStack* identified 60, approximately half as many. While *miRador* found the most known miRNAs, it also predicted far more miRNAs that are not known than the other two tools. In terms of precision, all tools had precision within 10% of one another, though *miRador* was the lowest at 0.785 while miRDeep-P2 was the highest at 0.874 (**Table 2**). We also explored the overlap between these predictions to determine if any of these prediction tools were encapsulated by one of the others. In the case of these *Arabidopsis* libraries, we found that only five of the total 82 miRNAs identified by *Shortstack* were exclusive to it (**Figure 2a**). While we did observe some overlap between the predictions made by both *miRador* and *miRDeep-P2*, 53.9% of *miRador*’s predictions were not observed by *miRDeep-P2*, and conversely 37.1% of *miRDeep-P2*’s predictions were not observed by *miRador*.

**Figure 2.**
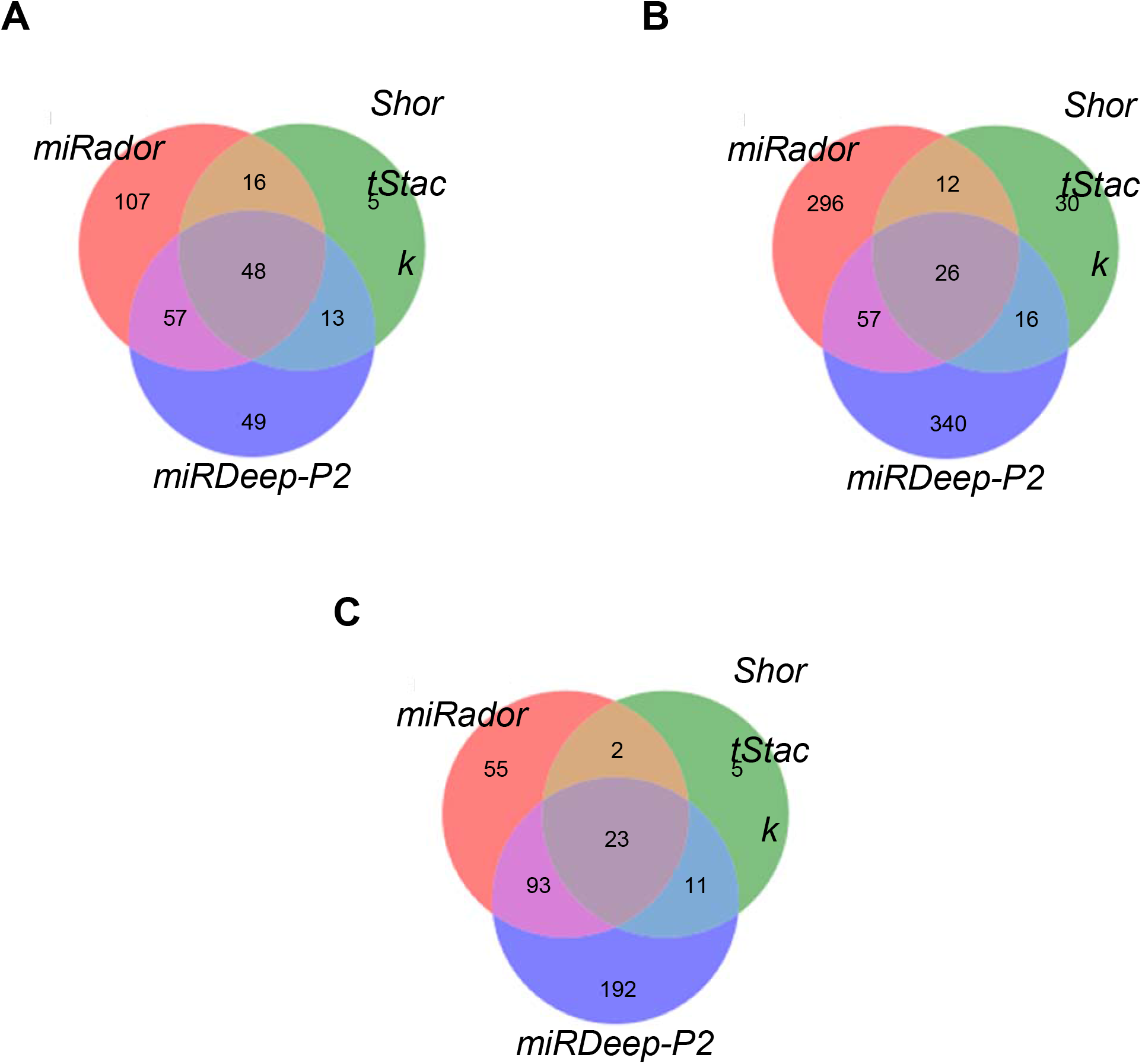
Overlap across the three prediction tools for miRNA predicted from three plant species. In each panel, the number of distinct candidate miRNAs that are found by each tool, and those predictions that were found in common by the different prediction tools, indicated as a Venn diagram. A. The number of distinct candidate *Arabidopsis* miRNAs that are found by each tool, and those predictions that were commonly found by the different prediction tools *Arabidopsis* miRNAs. B. The number of distinct candidate rice miRNAs that are found by each tool, and those predictions that were commonly found by the different prediction tools C. The number of distinct candidate maize miRNAs that are found by each tool, and those predictions that were commonly found by the different prediction tools.

Further predictions conducted in the 30 rice libraries resulted in slightly contrasting results to the case of *Arabidopsis*. Here, *miRDeep-P2* predicted the most (439) miRNAs, 178 of which were novel. *ShortStack* predicted the fewest miRNAs (84), while *miRador* predicted 357 miRNAs. In the rice libraries, we observed that each tool predicted several miRNAs that no other tool predicted (**Figure 2b**). We also found the precision of the three prediction tools was fairly comparable, this time only separated by 5% (**Table 3**). These findings were largely similar in maize where *miRDeep-P2* had the greatest precision at 0.887 and *ShortStack* had the smallest at 0.805 (**Figure 2c and Table 4**). Overall, these results indicate that each tool is comparable in terms of precision, though each tool uniquely identifies several miRNAs that the other tools do not.

Finally, we compared the runtimes of each tool with several *Arabidopsis*, rice, maize, and wheat libraries (**Figure 3**). We utilized *miRador* in its sequential run-mode to compare single core executions with *ShortStack* and *miRDeep-P2*. Of note, however, is that *ShortStack* is a tool that discovers and annotates small RNA clusters while also identifying *MIRNA* genes. Given this multi-functionality of *ShortStack*, we note that a comparison to its runtime is not necessarily 1-to-1. However, given that we have utilized *ShortStack* as a comparison of predictability and it is a commonly utilized tool for predicting miRNAs, we have opted to include its runtimes in our comparison.

**Figure 3:**
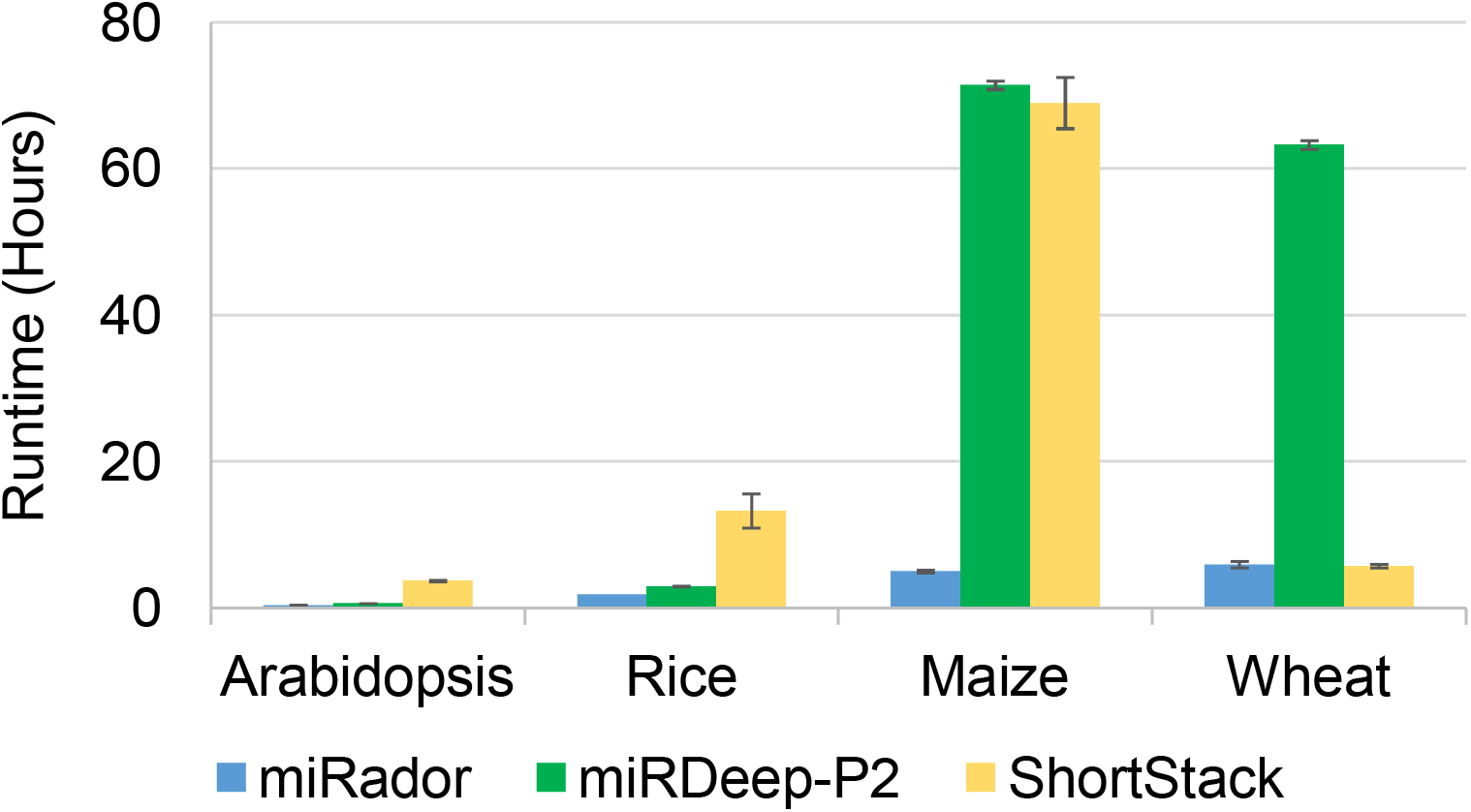
Runtime comparison of *miRador* to *miRDeep-P2* and *ShortStack* Comparison of execution times of *miRador, miRDeep-P2* and *ShortStack* across 21 *Arabidopsis* (116 Mb), 30 rice (364 Mb), 44 maize (2.1 Gb), and seven wheat (17 Gb) sRNA libraries. Execution was run using default parameters on the same server.

In our analysis of runtime performance, we predicted miRNAs in several libraries from four organisms: *Arabidopsis:* 116 MB (21 libraries), rice: 364 MB (30 libraries), maize: 2.1 GB (44 libraries) and wheat: 17 GB (7 libraries). Given that each tool utilizes *bowtie* to map libraries to the genome and both *ShortStack* and *miRDeep-P2* require bowtie indices to be created prior to execution, we did not include the optional *bowtie-build* step of *miRador* in these runtimes. In our assessment of *Arabidopsis*, rice, and maize libraries, we observed that *miRador* was the fastest tool. Notably, however, is the quick runtime of *ShortStack* in its prediction of wheat miRNAs relative to its runtime in the other organisms. *ShortStack*’s runtime scales with the number of libraries used as an input, due to its extensive effort in assigning weighted abundances to multi-mapped reads, a consideration not made by either *miRador* or *miRDeep-P2*. (Johnson et al., 2016). Given just seven libraries as an input, we found that this bottleneck was not as detrimental to its runtime leading *ShortStack* to be the fastest tool at predicting wheat miRNAs at 5.77 hours. *miRador* was only slightly slower at 5.97 hours while *miRDeep-P2* was the slowest at 63.27 hours. The runtime of *miRador* for predicting maize miRNAs was particularly strong, as its predictions across 44 sRNA libraries completed in an average of 5.05 hours, whereas both other tools took over three days to complete. Overall, we found that the runtime of *miRador* was, in our opinion, impressively quick. It outperformed the other tools in nearly every tested case, and in the case of wheat, it was nearly as fast as *ShortStack*. Its demonstrated scalability, supporting an ability to predict miRNAs in large genomes with many input libraries. Additionally, *miRador* has the ability to utilize multiple cores to improve prediction times, which can enable far larger analyses than those that we tested.

## DISCUSSION

In this paper, we describe *miRador*, a novel plant miRNA prediction tool that utilizes the most current community-developed plant miRNA annotation criteria. Unlike previous studies, we utilized PARE libraries to assess the quality of miRNA predictions beyond just the sequences that exist in miRBase, giving a better representation of the predictability of miRNA prediction tools. In addition to its strong predictive capabilities, we showed that *miRador* is faster than existing tools without compromising on predictive efficiency. Additionally, we developed an annotation function to annotate miRNAs with respect to their novelty, and similarity to known miRNAs, and we exported this function for use by other miRNA prediction tools.

In assessing the utility of *miRador* for miRNA predictions, we used PARE to identify the candidate miRNAs that have evidence of cleavage at their predicted targets, as identified by *sPARTA*. We utilized sRNA and PARE libraries from *Arabidopsis*, rice, and maize to identify the precision of *miRador, ShortStack*, and *miRDeep-P2*. We largely found that the precision of each prediction tool was comparable, though the number of miRNAs predicted by each tool varied by organism. *ShortStack* consistently identified the least miRNAs, though its predictions were not completely encapsulated by the predictions of the other two tools. *miRDeep-P2* and *miRador* had large overlaps in all organisms, but each tool also uniquely predicted numerous miRNAs. Overall, our findings suggest that each tool may be used to predict miRNAs with similar precision to one another, but none of these tools are all encapsulating, and thus there may be utility in running more than one tool when predicting miRNAs in a set of sRNA libraries.

In our attempts of validating candidate miRNA activity by target prediction and validation with PARE, we utilized PARE from corresponding tissues from which the sRNA data was acquired, although these were not always from the same plants or tissues. This may have resulted in reduced precision, particularly in the case of *Arabidopsis* in which most tools performed their worst in terms of precision. In the case of rice, however, we utilized well staged low-input sRNA and PARE sequencing libraries generated in triplicates from small amounts of tissue (Jiang et al., 2020). These tissue-specific datasets may provide cleaner results with less noise in the PARE data than what was found in the *Arabidopsis* and maize libraries resulting in the best predictions for all tools. In separate analyses not presented here, we found that rice miRNAs predicted and validated in inconsistent tissues had far less precision than those observed in the data presented in this study. In these analyses, the highest precision observed were the *miRador* predicted miRNAs with 60% had PARE evidence of cleavage at their predicted targets. This is in stark contrast to the 91% observed in the low-input sRNA and PARE datasets presented above. Such poor performance is likely attributable to the predicted miRNAs not accumulating in the tissues from which the PARE was acquired, thereby resulting in their classification as a false positive. Therefore, we note the importance of utilizing matching tissue types in sRNA and PARE datasets when validating predicted miRNAs.

Although we are confident in the quality of *miRador*, it is not without limitations. In its identification of candidate miRNA genes, *miRador* first searches for inverted repeats in a genome assembly using *einverted.* This worked well using high quality genome builds with which we performed the tests, but *miRador* may miss several miRNAs when predicting in novel genomes comprised of several disconnected scaffolds and contigs (an issue of diminishing concern as long-read sequencing and assemblies begin to predominate). We also largely utilized the community-established guidelines for plant miRNA annotations; these guidelines could be fine-tuned even further with machine learning to minimize false positives and false negatives. These additional methods would also allow for confidence scores to be assigned to the resulting predictions. We did acknowledge these limitations when building *miRador*, and thus it was largely developed with modular functions such that adjustments to its prediction filters could be established without overhauling the entire tool.

Despite these limitations, we assert that we have developed a tool that is as precise as other plant miRNA prediction tools while being far faster. *miRador* is highly scalable, ensuring its ability to predict miRNAs with large genomes with many sRNA libraries. The additional annotation component of *miRador*, which has been exported for use by other prediction tools, provides users great insights into the novelty of their predicted miRNAs. Altogether, *miRador* is a highly capable, standalone, plant miRNA prediction and annotation tool.

## MATERIALS AND METHODS

### Software and data availability

A description of this pipeline and overall algorithm are described in the Results section above. Further details are available in the README file which may be viewed on Github: https://github.com/rkweku/miRador.

We utilized public *Arabidopsis*, rice, maize, and wheat sRNA and PARE datasets for miRNA prediction and validation. These libraries, and their GEO accession numbers, are listed in **Supplementary Table 1**.

## ACKNOWLEDGEMENTS

We would like to thank both Joanna Friesner and Michael Axtell for comments on the manuscript. We thank members of the Meyers lab for helpful discussions. This work was supported by funding provided by the National Science Foundation award #1754097 to B.C.M, and resources provided by the Donald Danforth Plant Science Center and the University of Missouri - Columbia.

